# Computational genomic analysis of the lung tissue microenvironment in COVID-19 patients

**DOI:** 10.1101/2021.05.28.446250

**Authors:** Krithika Bhuvaneshwar, Subha Madhavan, Yuriy Gusev

## Abstract

The coronavirus disease 2019 (COVID-19) pandemic caused by the SARS-CoV-2 virus has affected over 170 million people, and caused over 3.5 million deaths throughout the world as of May 2021. Although over 150 million people around the world have recovered from this disease, the long term effects of the disease are still under study. A year after the start of the pandemic, data from COVID-19 recovered patients shows multiple organs affected with a broad spectrum of manifestations. Long term effects of SARS-CoV-2 infection includes fatigue, chest pain, cellular damage, and robust innate immune response with inflammatory cytokine production. More clinical studies and clinical trials are needed to not only document, but also to understand and determine the factors that predispose certain people to the long term side effects of his infection.

In this manuscript, our goal was to explore the multidimensional landscape of infected lung tissue microenvironment to better understand complex interactions between SARS-CoV-2 viral infection, immune response and the lungs microbiome of COVID-19 patients. Each sample was analyzed with several machine learning tools allowing simultaneous detection and quantification of viral RNA amount at genome and gene level; human gene expression and fractions of major types of immune cells, as well as metagenomic analysis of bacterial and viral abundance. To contrast and compare specific viral response to SARS-COV-2 we have analyzed deep sequencing data from additional cohort of patients infected with NL63 strain of corona virus.

Our correlation analysis of three types of measurements in patients i.e. fraction of viral RNA (at genome and gene level), Human RNA (transcripts and gene level) and bacterial RNA (metagenomic analysis), showed significant correlation between viral load as well as level of specific viral gene expression with the fractions of immune cells present in lung lavage as well as with abundance of major fractions of lung microbiome in COVID-19 patients.

Our exploratory study has provided novel insights into complex regulatory signaling interactions and correlative patterns between the viral infection, inhibition of innate and adaptive immune response as well as microbiome landscape of the lung tissue. These initial findings could provide better understanding of the diverse dynamics of immune response and the side effects of the SARS-CoV-2 infection.

## INTRODUCTION

The coronavirus disease 2019 (COVID-19) pandemic caused by the SARS-CoV-2 virus has affected over 170 million people, and caused more than 3.5 million deaths throughout the world as of May 2021 [1]. Majority of the patients infected were reported to have mild disease (about 75-80%), about 15-20% of patients need hospitalization, and about 5-10% need critical care [2, 3].

Scientists have been studying the molecular underpinnings of this infection. Several large multi-institutional efforts have been started with the goal of understanding the underlying mechanisms of this infection. One such effort is the COVID Human Genetic Effort which explores the genetic and immunological reasons for the various clinical severities of this disease [4, 5]. Some reports found that patients who get severe disease lacked effective immune response due to either a mutation or a lack of effective viral response to overcome severe disease [6]. This effort also highlighted that inborn errors of type I interferons (IFN), or auto-antibodies to type I interferons associated with COVID-19 based pneumonia [6], and its therapeutic applications [7]. Another effort is the National COVID Cohort Collaborative, or the N3C organized by the NIH National Center for Advancing Translational Sciences (NCATS), a large effort for collecting and harmonizing data derived from electronic health records (EHRs) from different institutions for collaborative research [8]. Other large efforts include the COVID Symptom Study, an app for tracking symptoms [9]; and NIH COVID digital pathology repository that collects whole slide images (WSI) of COVID related pathology [10]. The National Heart, Lung, and Blood Institute (NHLBI) is conducting studies to test if medications are safe and can help individuals recover from COVID-19 [11]. Another NIH-funded study called the Collaborative Cohort of Cohorts for COVID-19 Research (C4R) Study is working to use population data determine factors that predict disease severity and long-term health impacts of COVID-19 [12, 13].

Although over 150 million people around the world have recovered from this disease, the long term effects of the disease are still under study. A year after the start of the pandemic, data from COVID-19 recovered patients shows multiple organs affected with a broad spectrum of manifestations. Long term effects of SARS-CoV-2 infection, also known as ‘long COVID’ includes fatigue, chest pain, cellular damage, and robust innate immune response with inflammatory cytokine production [2, 14–16]. These collection of side effects are referred to as post-acute sequelae of SARS-CoV-2 infection (PASC) [17]. In Feb 2021, NIH announced an initiative to invest $1.15 billion over four years to fund studies to build a biospecimen bank and track recovery from long COVID referred to as the NIH PASC Initiative [18]. The UK National Institute for Health Research has also announced an investment of £18.5 million into the launch of four long COVID studies [19].

Such studies would enable understand this disease from a molecular perspective with the ultimate goal of finding better treatments. Researchers would be to use the big data collected from one of these large initiatives to build learning based models and determine the factors that predispose certain people to the long term side effects of this infection.

In this manuscript, our goal was to explore the multidimensional landscape of infected lung tissue microenvironment to better understand complex interactions between SARS-CoV-2 viral infection, immune response and the lungs microbiome of COVID-19 patients.

We report genomic analysis of deep sequencing data from publicly available RNA samples of lung lavage of COVID-19 patients. Each sample was analyzed with three different machine learning tools allowing simultaneous detection and quantification of viral RNA amount at genome and gene level; human gene expression and fractions of major types of infiltrating immune cells, as well as metagenomic analysis of bacterial and viral abundance in lower respiratory tract.

We also compared and contrasted specific immune responses to SARS-CoV-2 with the immune response to an infection by different coronavirus NL63. The NL63 coronavirus (HCoV-NL63) causes severe lower respiratory tract infection, and bronchiolitis disease in children, the elderly and the immunocompromised [20]. To conduct this comparison we have applied the same computational tools and analyzed deep sequencing data from an additional cohort of patients infected with NL63 strain of coronavirus.

## MATERIALS AND METHODS

Our goal was to explore the multidimensional landscape of infected lung tissue microenvironment to better understand complex interactions between virus, immune response and microbiome in the lungs of COVID-19 patients. By utilizing three types of machine learning based bioinformatics tools, we were able to detect and quantitate three different fractions of short reads from RNAseq data files: fraction of viral RNA (at genome and gene level), Human RNA (transcripts and gene level) and bacterial RNA (metagenomic analysis).

### Datasets

For this paper, we applied our analysis workflow on two datasets described. One of our goals was to explore sequences of the SARS-CoV-2 viral genome and gene level data and compare to other coronavirus genomes. We chose a Human NL63 coronavirus (HCoV-NL63) dataset as this virus is known to affect children, the elderly and the immunocompromised [20]. This virus is one of the many human endemic coronaviruses (hCoV) that include HCoV-229E, and NL63-CoV that are from *alphacoronavirus* group; and OC43 and HKU1 that are from the *betacoronavirus* group [21]. This dataset allowed for direct comparison with SARS-CoV-2 dataset and has been presented in the main body of the manuscript.

a. ***SARS-CoV-2* Dataset:** We downloaded RNA-seq data from 5 patients affected with the SARS COV 2 virus from the NCBI SRA data repository PRJNA605983 (SRP249613) [22, 23]. These 5 patients were from the early stage of the Wuhan seafood market pneumonia virus outbreak in China. The downloaded data were raw sequences in the form of .FASTQ files. Total RNA was extracted from bronchoalveolar lavage fluid and then next generation sequencing (NGS) was performed using the Illumina platform. Out of the 5 samples, 4 were profiled on the Illumina HiSeq 3000 platform, and one on the Illumina HiSeq 1000 platform.
b. **Human NL63 coronavirus dataset (referred to as the *NL63-CoV* dataset in this paper):** We found a public dataset was from 5 pediatric patients with severe lower respiratory infection by NL63 coronavirus with deep sequencing data performed on Illumina HiSeq platform. The downloaded data were raw sequences in the form of .FASTQ files obtained from NCBI PRJA601331 [24, 25]. In this dataset, nasopharyngeal swab samples were obtained from children with severe acute respiratory infection (SIRS) and then sequenced using the Illumina HiSeq 2500 platform. The researchers who submitted the dataset noted in their manuscript (Zhang et al [25]) that only partial genome sequences were obtained by NGS methods from the 5 samples [25].

## METHODS

Our analysis pipeline is shown in Figure 1. Our analysis workflow includes three main sections: analysis and exploration of the viral, immune, and bacterial environment in the RNA-seq data.

**Figure 1.**
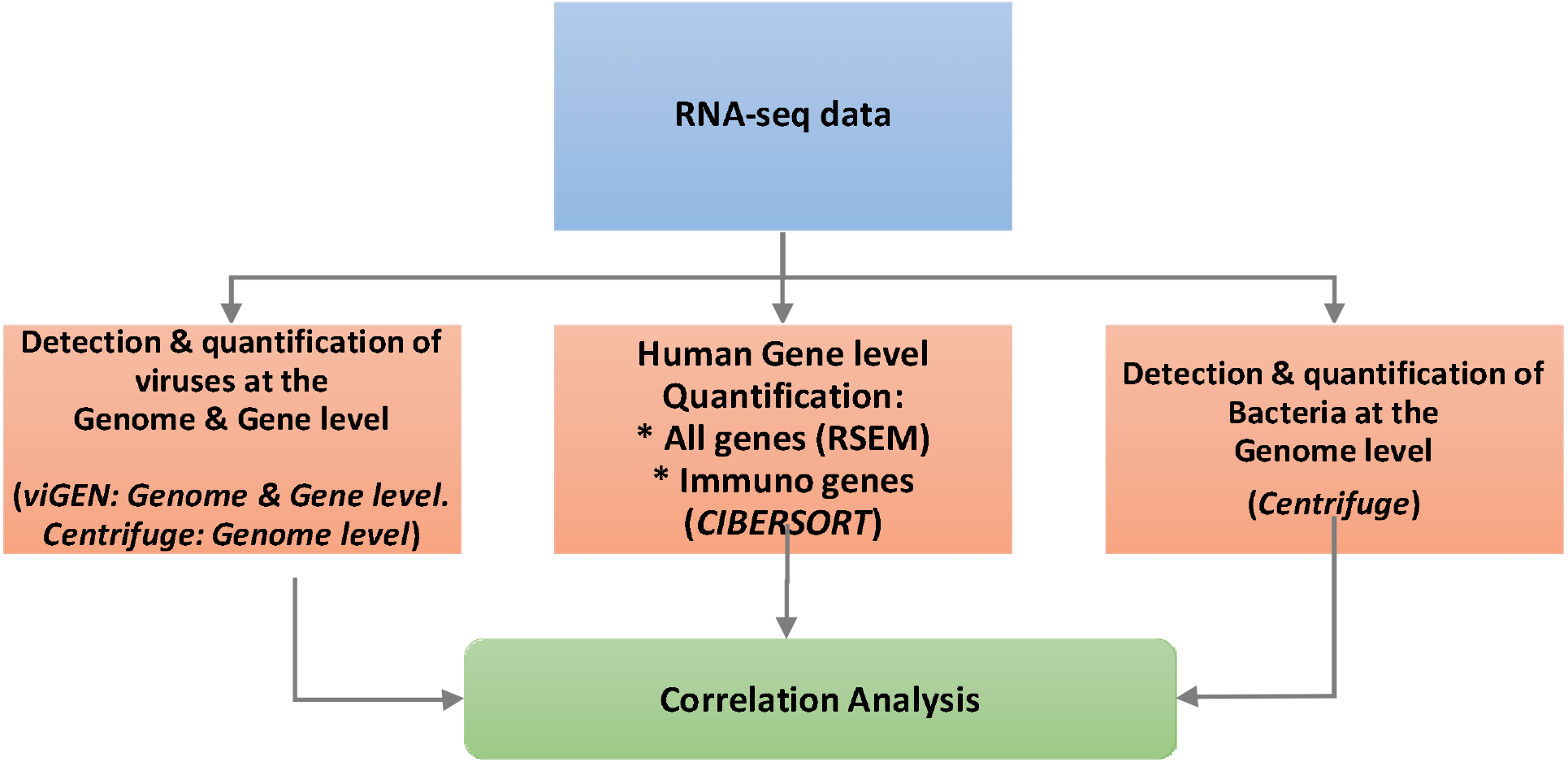
Data Analysis Workflow.

### Analysis of the viral environment in RNA-seq data

We first compiled our viral reference genome by downloading the viral sequences with humans as host from the NCBI collection of viral genomes [26]. We then ran our viGEN viroinformatics pipeline [27] on the samples from the *SARS-CoV-2 and the NL63-CoV datasets*. viGEN is an open source bioinformatics pipeline which allows for not only the detection and quantification of viral RNA, but also variants in the viral transcripts [27]. For this paper, we applied the viGEN pipeline that uses Bowtie2 aligner [28] to find the viruses detected in the two datasets. Bowtie2 is an alignment algorithm that uses a combination of index–assisted seed alignment and single-instruction multiple-data (SIMD) based dynamic parallel processing to achieve fast, accurate and sensitive alignment of sequencing data [28]. We also corroborated these results with the help of an additional pipeline CENTRIFUGE [29] to detect and quantitate abundance of viral species. Then we applied our quantification algorithm of viral RNA at the gene/CDS level, which is part of the viGEN pipeline [27]. This produced gene counts of viral RNA on the input datasets.

### Analysis of the immune environment in the RNA-seq data

We applied our immuno-genomics pipeline to the *SARS-COV-2 and NL63-CoV datasets*. The raw sequences were first aligned to the human reference genome version hg38 using an open source RNA-seq alignment and quantification pipeline that used the RSEM-STAR aligner algorithm [30]. This analysis was performed in the Seven Bridges (SB) Cancer Genomics Cloud (CGC) Platform [31, 32]. The output of this pipeline were gene and isoform level data in the form of both raw counts and transcripts per million (TPM) values. Out of the ~20,000 genes in the human genome, the gene quantification data in the form of TPM values were extracted for a subset of 530 immune related genes. This gene matrix was input into a public online tool CIBERSORT [33]. CIBERSORT is a virtual flow cytometry tool that estimates the abundance of immune cell infiltrates in a mixed cell population using gene expression or RNA-seq data. It is machine learning algorithm that uses nu-linear support vector regression (ν-SVR) to perform deconvolution of the input mixture. ν-SVR is a type of support vector machine (SVM) wherein a hyperplane that maximally separates both classes is discovered [34]. The CIBERSORT analysis was performed with quantile normalization disabled and a permuted 500 times. The output of CIBERSORT were in the form of fractions of 22 immune cell infiltrates across the input samples [34].

### Analysis of the bacterial environment in the RNA-seq data

Additional analysis was performed on the same RNA samples by applying metagenomics pipeline CENTRIFUGE [29] to detect and quantitate abundance of bacterial species comprising lung microbiome. Centrifuge is a machine learning algorithm that uses an indexing scheme based on the Burrows-Wheeler transform (BWT) and the Ferragina-Manzini (FM) index. It was designed and optimized specifically for metagenomics applications [29]. We ran the Centrifuge analysis pipeline on the Seven Bridges CGC platform [31, 32] on the input same FASTQ files from the datasets. The index file containing the reference genomes for human, prokaryotic genomes, and viral genomes including 106 SARS-CoV-2 complete genomes was provided by the CENTRIFUGE team on their website [35]. We also used the online web application Pavian [36, 37] for visual analysis of results generated with Centrifuge.

### Correlation analysis

Once the three types of results were obtained for each sample in the datasets, downstream correlation analysis was performed. This allowed us to explore possible interactions and regulation of viral infection with local immunological environment in lungs of infected patients, as well as, microbiome profiles measured by changes in abundance of multiple bacterial species in the lung. The Pearson correlation analysis was performed in the R statistical software [38].

## RESULTS

Our goal was to explore multidimensional landscape of infected lung tissue microenvironment to better understand complex interactions between virus, immune response and microbiome in the lungs of COVID19 patients. By utilizing three types of bioinformatics workflows and tools, we were able to detect and quantitate three different fractions of short reads from RNAseq data files: fraction of viral RNA (at genome and gene level), Human RNA (transcripts and gene level) and bacterial RNA (metagenomic analysis). Correlation analysis of these three types of measurements in patients has showed significant correlation between viral load as well as level of specific viral gene expression with the fractions of immune cells present in lung lavage as well as with abundance of major fractions of lung microbiome.

### Section 1: Results of analysis on the SARS-CoV-2 dataset

#### 1.1 Results of analysis of the viral environment

Supplementary File 1A shows the estimated copy number of viral genomes detected in lung lavage samples of the *SARS-CoV-2 dataset* obtained using the viGEN pipeline. We can clearly see the presence of the various strains SARS-CoV-2 virus in all of the 5 patients in this dataset. The SARS-CoV-2 virus in these samples has also been detected and corroborated using the CENTRIFUGE metagenomics pipeline (Supplementary File 1B).

Supplementary File 1C shows the estimated level of viral gene expression counts in the patients from the *SARS-CoV-2 dataset*. The three prime UTR region in the region ranging from nucleotide position 29675 to 29903 shows one of the largest abundance of viral gene expression with raw gene counts more than 1000 for most patients. We also see the presence of the older SARS coronavirus (SARS-CoV) which was prevalent in Asia in 2003 [39]. This makes sense as these RNA-seq samples were from patients China. As expected, we see presence of flu and common cold viruses including Human Coronavirus 229E and Influenza A virus (A/Shanghai/02/2013(H7N9)).

#### 1.2 Results of analysis of the immune environment

Next, we assessed the immuno-profile of samples from the *SARS-CoV-2 dataset* (Figure 2A). It shows a summary of the estimated fractions of 22 types of immune cells detected in lung lavage and indicates Macrophages M2, T cells CD4 naïve, Natural Killer (NK) cells resting and Monocytes as the most abundant types of immune cells in this dataset.

**Figure 2.**
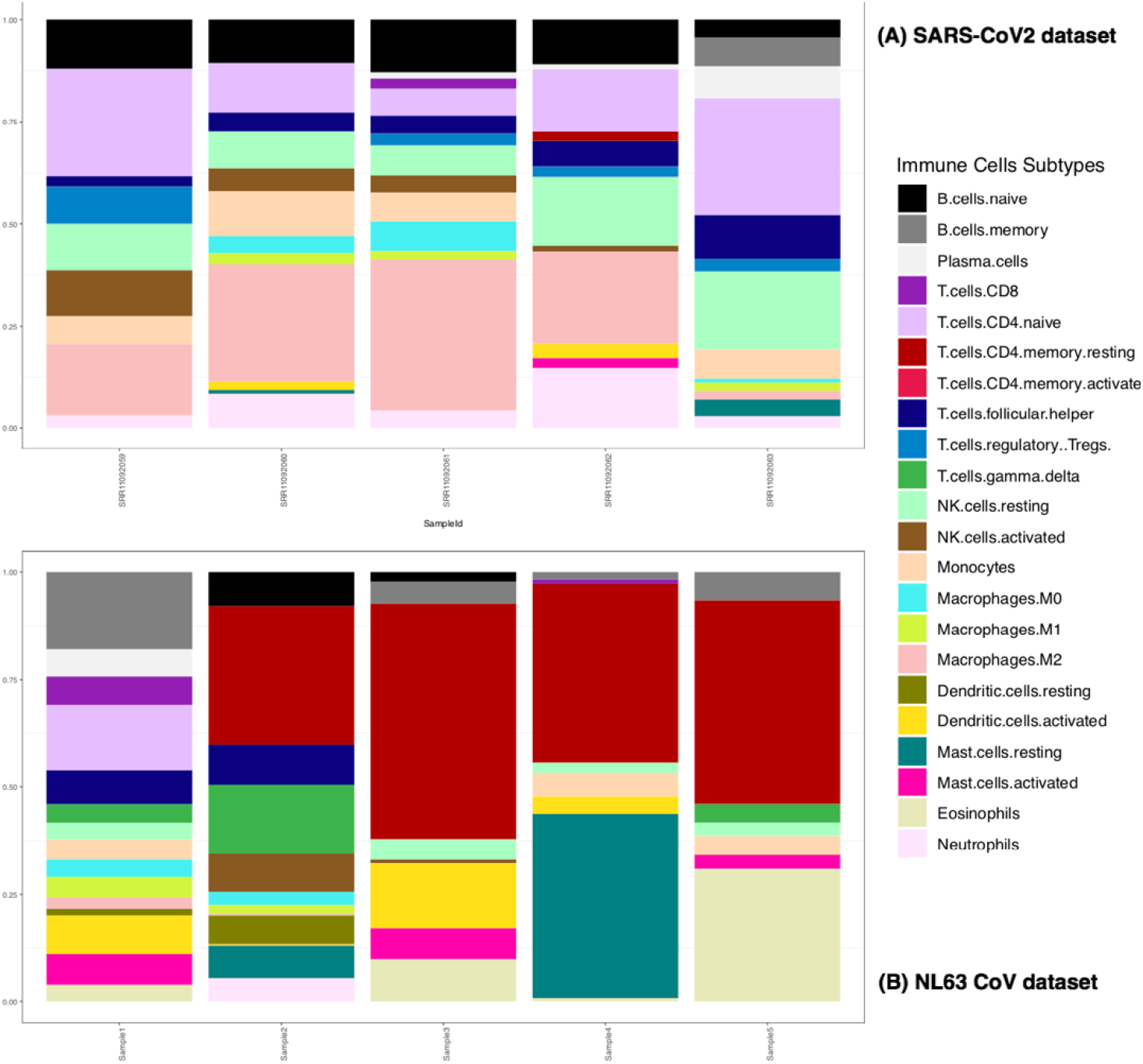
CIBERSORT output : estimated fractions of 22 types of immune cells detected in lung lavage of (A) 5 patients from the SARS-COV-2 dataset and (B) 5 patients from the NL63-COV-Hiseq dataset. There is one stacked bar per patient.

#### 1.3 Results of analysis of the bacterial environment

The metagenomic analysis of the patients in the *SARS-CoV-2 dataset* (Table 1) shows the abundance of the top 20 bacterial species in the lung microbiome. Figure 3 shows a Sankey diagram visualization of the bacterial species. A Sankey diagram [40] is a flow diagram in which the width of the arrows is proportional to the abundance (i.e. number of reads) of the bacterial species, which are collated together by the taxonomy of bacteria. Hence the most abundant bacterial species will be shown on the right most side with the largest width. The Sankey diagrams indicated bacterial species *Clostridium botulinum* and *Clostridium tetani* to be one of the most abundant species in these samples

**Table 1.** Abundance of top bacterial species in lung microbiome of patients in the SARS-CoV-2 dataset. Showing top 20 sorted based on minimum counts

| Name | Minimum | SRR11092059 | SRR11092060 | SRR11092061 | SRR11092062 | SRR11092063 |
| --- | --- | --- | --- | --- | --- | --- |
| *Clostridium tetani* | 257477 | 886195 | 305193 | 257477 | 433532 | 355297 |
| *Clostridium botulinum* | 218725 | 1499933 | 252669 | 218725 | 317989 | 264103 |
| *Trichormus azollae* | 29537 | 31171 | 37568 | 29537 | 57222 | 46985 |
| *Spirosoma pollinicola* | 19190 | 246664 | 24916 | 25987 | 24161 | 19190 |
| *Prevotella oris* | 9083 | 9757 | 9986 | 9083 | 13242 | 10862 |
| *Staphylococcus aureus* | 8482 | 1142840 | 46143 | 28001 | 13914 | 8482 |
| *Stenotrophomonas maltophilia* group | 7784 | 1312705 | 116353 | 116221 | 24795 | 7784 |
| *Stenotrophomonas maltophilia* | 7784 | 1312705 | 116353 | 116221 | 24795 | 7784 |
| *Prevotella denticola* | 7630 | 7955 | 8340 | 7630 | 10548 | 8846 |
| *Prevotella fusca* | 7036 | 7530 | 7895 | 7036 | 10089 | 8459 |
| *Roseimicrobium* sp. ORNL1 | 6499 | 6872 | 8053 | 6499 | 12297 | 10272 |
| *Enterobacter cloacae* complex | 6377 | 698959 | 68035 | 67773 | 13735 | 6377 |
| *Prevotella ruminicola* | 5829 | 6073 | 6150 | 5829 | 8186 | 6958 |
| *Enterobacter kobei* | 4130 | 452826 | 44783 | 44580 | 8897 | 4130 |
| *Pseudomonas putida* group | 4105 | 556693 | 51160 | 51556 | 11347 | 4105 |
| *Pseudomonas putida* | 3667 | 516644 | 47579 | 47876 | 10105 | 3667 |
| *Sphingomonas paucimobilis* | 1766 | 296679 | 1766 | 23579 | 13965 | 2448 |
| *Enterobacter cloacae* | 1461 | 160877 | 15373 | 15426 | 3054 | 1461 |
| *Lacrimispora saccharolytica* | 1140 | 159024 | 13606 | 13991 | 2903 | 1140 |
| **Total Bacterial reads in the sample** | **984975** | **10726961** | **1358144** | **1328864** | **1353674** | **984975** |
| **Total Microbial reads in the sample** | **1016348** | **10849767** | **1412506** | **1578271** | **1436901** | **1016348** |

**Figure 3.**
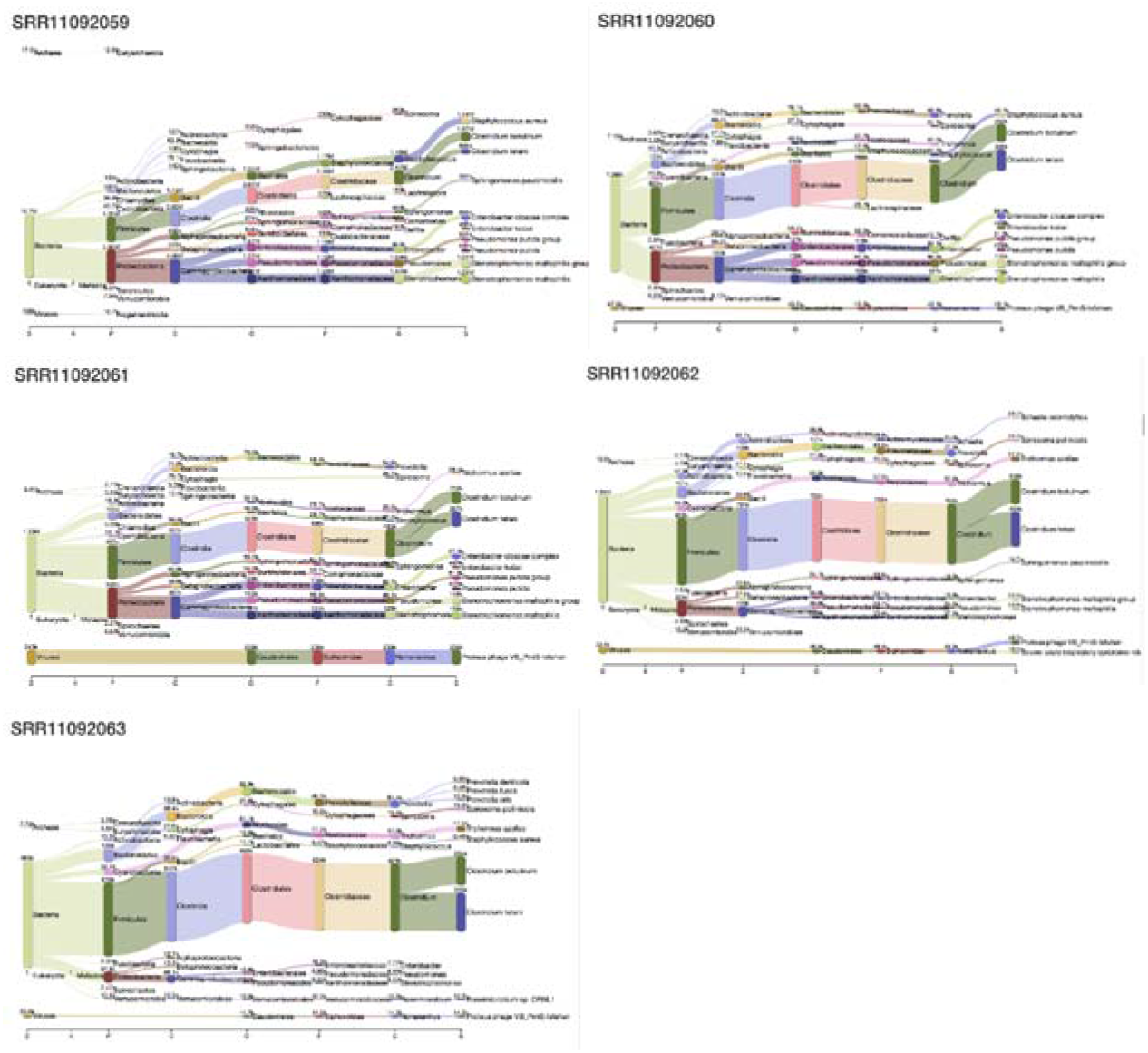
Lung microbiome profile for the SARS-CoV-2 dataset represented as a Sankey diagram visualization of the bacterial species.

#### 1.4 Results of correlation between genomic and immunological data

We found significant correlation between viral load (genomic and gene level) and immune-profile of the patients in the *SARS-CoV-2 dataset.*

Figure 4 (A and B) shows the summary of the statistically significant correlations between viral gene expression and immunological cell types for the SARS-CoV-2 dataset. The full correlation matrix is provided as Supplementary File 2A (genome level) and Supplementary File 2B (gene level). Figure 4A shows the statistically significant correlation results between viral load (genome level) and fraction of immune cells. Results show that Macrophages M2 are positively correlated with the SARS-CoV-2 virus genome counts, while NK cells activated, monocytes and T cells CD4 Naïve were negatively correlated with genome counts. Figure 4B represents statistically significant correlations between viral gene expression (gene level) and fraction of immune cells. Activated NK cells and monocytes were found to be inversely correlated with the gene counts of the 3 prime and 5 prime UTR regions respectively. Monocytes were also found to be inversely correlated with other regions of the SARS-CoV-2 virus genome including membrane glycoprotein, envelope protein, nucleocapsid phosphoprotein and more.

**Figure 4:**
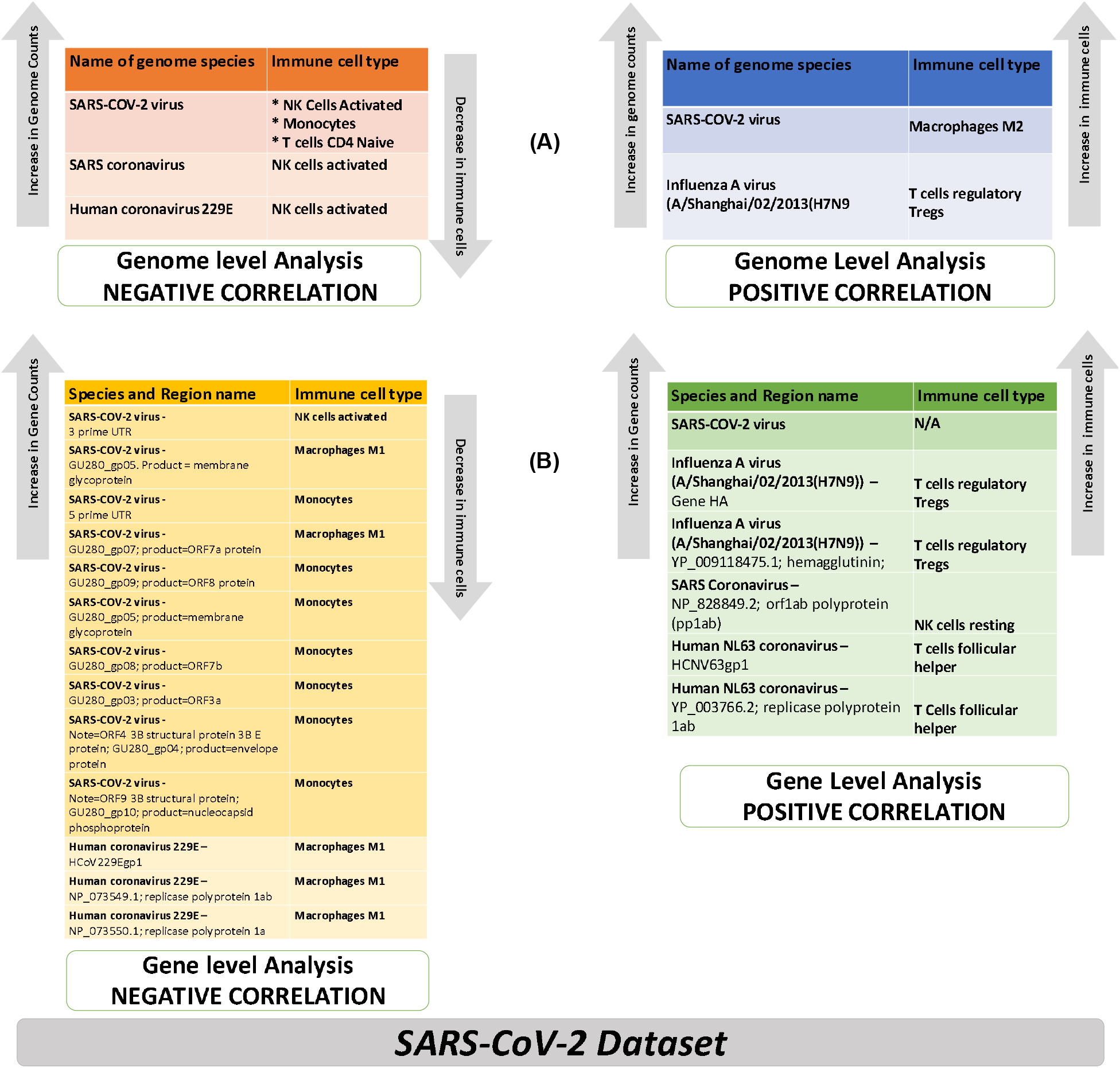
Summary of the statistically significant correlations between viral gene expression and immunological cell types for the SARS-CoV-2 dataset. (A) Viral genome level correlation between. viral load and fraction of immune cells (B). Viral gene level correlation between viral gene expression and fraction of immune cells

#### 1.5 Results of correlation between bacterial abundance with immunological cell types

We chose a p-value cut off of 0.005 to get a short list of 70 statistically significant correlation results. Out of these 70 short listed results, only one was negatively correlated, while the rest of the 69 results were positively correlated. Monocytes were negatively correlated with *Schaalia odontolytica* bacterial species. NK cells activated and Eosinophils were positively correlated with the following families including *Citrobacter, Clostridium, Delftia, Enterobacter, Lactobacillus, Paenibacillus, Phyllobacterium and Pseudomonas, Staphylococcus and Stenotrophomonas* (Figure 5). The complete correlation results for this correlation analysis is available as Supplementary File 2C

**Figure 5:**
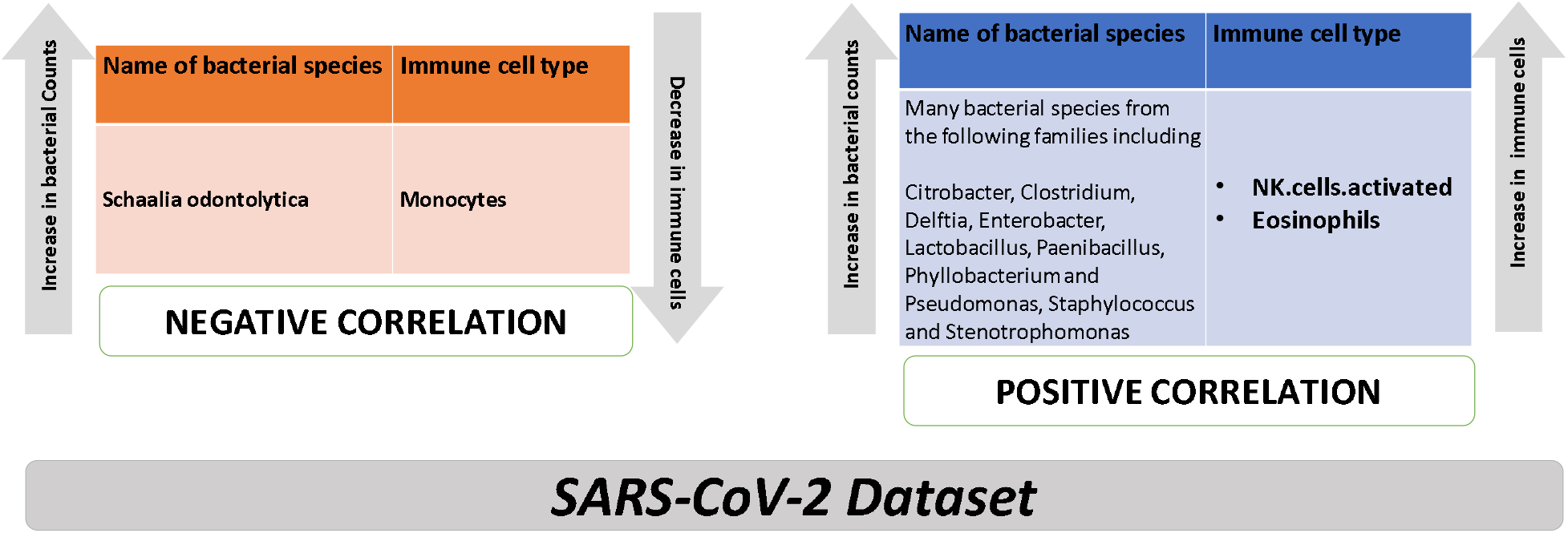
Summary of correlation analysis between bacterial abundance with immunological cell types for the SARS-CoV-2 dataset.

#### 1.6 Results of Correlation between viral load (genomic level) with bacterial abundance

We chose a p-value cut off of 0.005 to get a short list of 261 statistically significant features that were correlated between viral load (genomic level) and bacterial abundance. Due to the large number of results, we focused on the correlation results in the Severe acute respiratory syndrome coronavirus 2 (SARS-CoV-2) genome The complete correlation results for this correlation analysis is available as Supplementary File 2D. Of the 261 statistically significant features, only one bacterial species *Schaalia odontolytica* was positively correlated with the SARS-CoV-2 genome. The rest of the results were negatively correlated with the SARS-CoV-2 genome including *Citrobacter, Enterobacter, Lactobacillus, Paenibacillus, and Pseudomonas*. These results (summarized in Figure 6) are very similar to the correlation analysis of the bacterial abundance with immunological cell types for the SARS-CoV-2 dataset (Figure 5).

**Figure 6:**
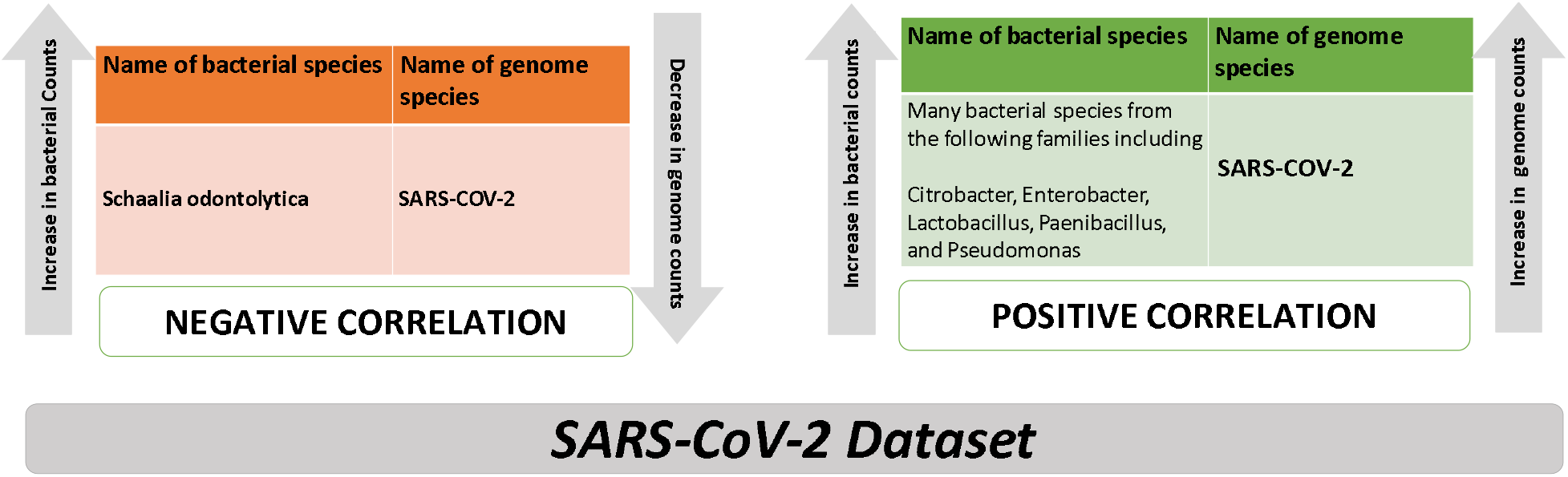
Correlation analysis between viral load (genomic level) with bacterial abundance in the SARS-CoV-2 dataset.

### Section 2: Results of analysis on the NL63-CoV dataset

#### 2.1 Results of analysis of the viral environment

Supplementary File 3A shows the estimated viral genome copy numbers in 5 pediatric patients from the *NL63-CoV dataset* obtained using the viGEN pipeline. Human coronavirus NL63 virus was detected in only two of the five pediatric patients. This same result is corroborated using the Centrifuge pipeline (Supplementary File 3B) which showed a read count of more than 100 for the same two samples. The researchers who submitted the dataset noted in their manuscript (Zhang et al [25]) that only partial genome sequences were obtained by NGS methods from the 5 samples [25]. We think this may have caused the non-detection of the NL63-CoV virus in some samples.

Supplementary File 3C shows the NL63 viral gene/CDS counts in the *NL63-CoV dataset.* Some of the most abundant regions include the CDS regions in the HCNV63gp1 and HCNV63gp2 gene that code for the *replicase polyprotein 1ab.* Other dominant regions include a CDS region in HCNV63gp2 gene that codes for a spike protein; and a CDS region in HCNV63gp4 gene that codes for a envelope protein (small membrane protein).

#### 2.2 Results of analysis of the immune environment

Next, we assessed the immuno-profile of samples from the *NL63-CoV dataset* (Figure 2B). It indicates T cells CD4 memory as the dominant immune cells.

#### 2.3 Results of analysis of the bacterial environment

The metagenomic analysis of the patients in the *NL63-CoV dataset* (Table 2) shows the abundance of the top 20 bacterial species in the nasopharyngeal microenvironment shows *Streptococcus pneumoniae* as one the dominant species

**Table 2.** List of top bacterial species dominating nasopharyngeal microenvironment microbiome of 5 NL63 patients based on maximum abundance. Showing top 20 sorted based on minimum counts

| Name | Minimum | Max | Sample1 | Sample2 | Sample3 | Sample4 | Sample5 |
| --- | --- | --- | --- | --- | --- | --- | --- |
| Streptococcus pneumoniae | 5121 | 196870 | 14355 | 39493 | 196870 | 5121 | 12381 |
| Streptococcus mitis | 4742 | 465843 | 28008 | 74808 | 465843 | 4742 | 20932 |
| Campylobacter concisus | 3187 | 536664 | 12525 | 536664 | 50631 | 3255 | 3187 |
| Prevotella melaninogenica | 2604 | 1058483 | 1058483 | 587266 | 395090 | 2604 | 13004 |
| Haemophilus influenzae | 2408 | 33233 | 10396 | 15293 | 33233 | 2408 | 23473 |
| Streptococcus pseudopneumoniae | 2265 | 72017 | 4201 | 9186 | 72017 | 2265 | 4597 |
| Prevotella jejuni | 1770 | 818696 | 41903 | 552126 | 818696 | 2692 | 1770 |
| Haemophilus haemolyticus | 1710 | 44072 | 14249 | 23371 | 31369 | 1710 | 44072 |
| Veillonella dispar | 1709 | 360869 | 54288 | 360869 | 347922 | 1709 | 3706 |
| Streptococcus salivarius | 1381 | 103093 | 1650 | 33522 | 103093 | 1381 | 3776 |
| Veillonella atypica | 1169 | 470491 | 132008 | 470491 | 392930 | 1878 | 1169 |
| Neisseria meningitidis | 1083 | 23338 | 12358 | 23338 | 4311 | 1083 | 2798 |
| Neisseria flavescens | 908 | 293417 | 293417 | 46429 | 12337 | 1595 | 908 |
| Streptococcus gwangjuense | 805 | 83898 | 5497 | 17502 | 83898 | 805 | 3638 |
| Streptococcus sp. 116-D4 | 637 | 48931 | 3913 | 16995 | 48931 | 637 | 2940 |
| Haemophilus sp. oral taxon 036 | 610 | 9971 | 2992 | 4245 | 9971 | 610 | 8280 |
| Fusobacterium pseudoperiodonticum | 530 | 3451701 | 3451701 | 1120955 | 27430 | 530 | 596 |
| Prevotella intermedia | 513 | 268641 | 165720 | 268641 | 99410 | 513 | 1128 |
| Streptococcus oralis | 502 | 82478 | 11027 | 81068 | 82478 | 502 | 2354 |

Supplementary File 4 shows the nasopharyngeal microbiome profile of pediatric patients from the *NL63-CoV dataset* represented as a Sankey diagram visualization of the bacterial species.

#### 2.4 Results of correlation between genomic and immunological data

We did not find many significant correlations between viral load and viral gene expression and immune-profile of the patients in the *NL63-CoV dataset.* This may have been attributed to the challenges the owners of this dataset faced with regards to partial genome sequences obtained during sequencing by NGS methods. While there were no significant correlation between viral load and viral gene expression and immune-profile for the NL63 coronavirus species, we did find some significant correlation with other similar coronavirus species, the Human Coronavirus 229E. Figure 5 shows the summary of the statistically significant correlations between viral gene expression and immunological cell types for the *NL63-CoV dataset*. The full correlation matrix is provided as Supplementary File 5A (genome level) and Supplementary File 5B (gene level). Figure 5A represents the correlation at the genome level between viral load and fraction of immune cells. Figure 5B represents the gene level correlation between viral gene expression and fraction of immune cells.

#### 2.5 Correlation between metagenomic bacterial abundance with immunological cell types

We chose a p-value cut off of 0.005 to get a short list of 17 statistically significant correlation results. Out of these 17 short listed results, 16 were negatively correlated, and one positively correlated. B cells naïve were positively correlated with bacterial species *Mycoplasma orale.* Monocytes and Mast Cells resting were negatively correlated with many bacterial species from the following families including *Prevotella, Streptococcus and Veillonella* (Figure 6). The full correlation matrix results for this analysis is provided as Supplementary File 5B

#### 2.6 Correlation between viral load (genomic level) with metagenomic abundance

We chose a P value cut off of 0.05 to get a list of 15 statistically significant features that correlated viral load (genomic level) with metagenomic abundance. We focused on the results in the NL63 coronavirus genome. We found no bacterial species positively correlated with the NL63 coronavirus genome. There were a few bacterial species negatively correlated with the NL63 coronavirus genome including *Fusobacterium, Prevotella, Streptococcus, Treponema and Veillonella.* The full correlation matrix results for this analysis is provided as Supplementary File 5D.

## DISCUSSION

Our goal was to explore multidimensional landscape of infected lung tissue microenvironment to better understand complex interactions between virus, immune response and microbiome in the lungs of COVID-19 patients in comparison with NL63-CoV patients. By utilizing three types of machine learning based bioinformatics tools, we were able to detect and quantitate three different fractions of short reads from RNAseq data files: fraction of viral RNA (at genome and gene level), Human RNA (transcripts and gene level) and bacterial RNA (metagenomic analysis).

### Immune landscape and correlation analysis in the *SARS-CoV-2* dataset

We compared the immune landscapes of the patients in the *SARS-CoV-2 and NL63-CoV* datasets with the help of box plots (Figure 9). Figure 9 indicates that Macrophages M2, T cells CD4 naïve, and Natural Killer (NK) cells are the most abundant in the immune landscape of the *SARS-CoV-2* dataset. On the other hand, T cells CD4 memory resting, and mast cells are the most abundant in the immune landscape of the *NL63-CoV* dataset.

**Figure 7:**
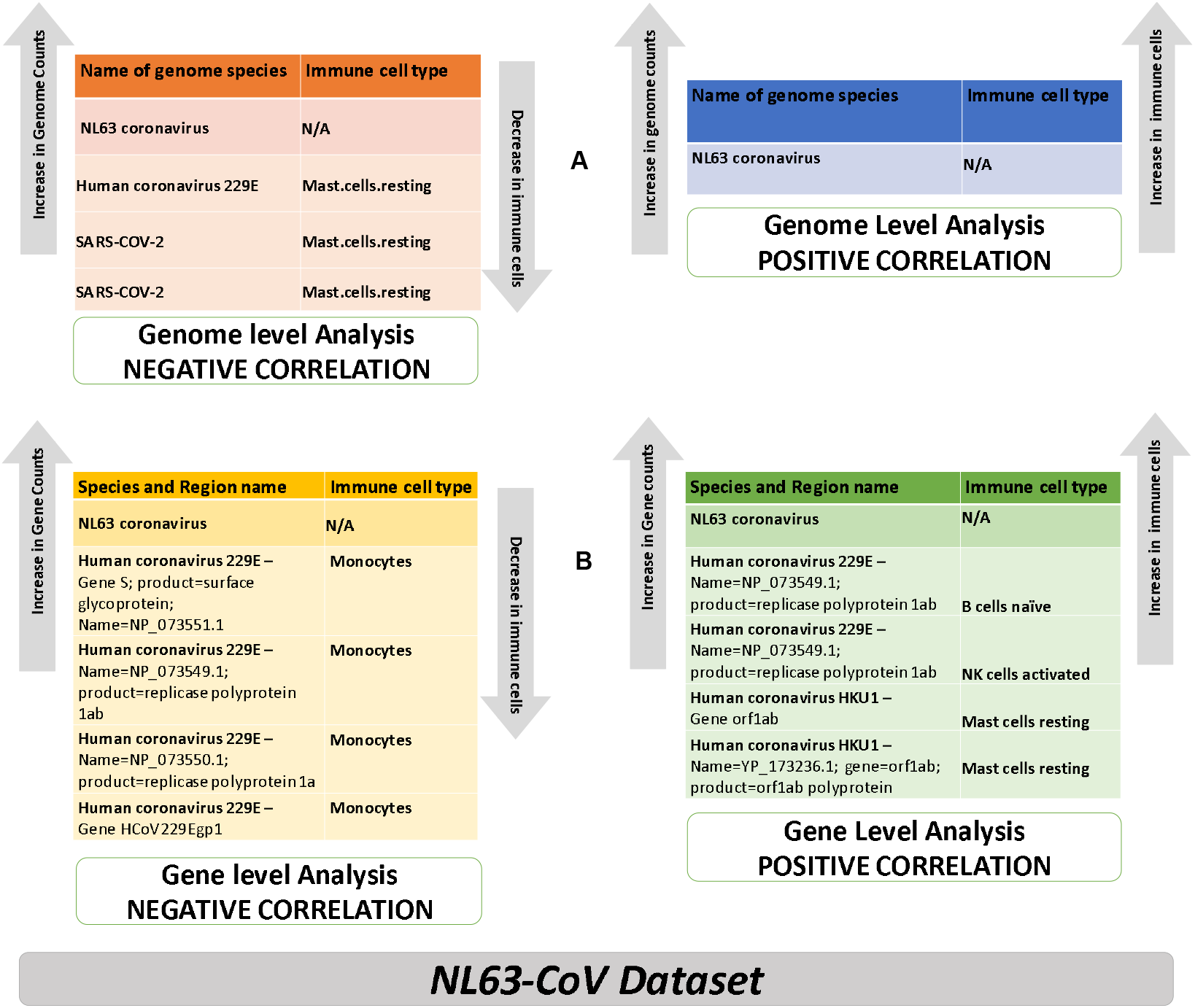
Summary of Statistically significant correlation between viral gene expression and immunological cell types for 4 NL63-CoV patients. A Viral genome level correlation (viral load vs. fraction of immune cells) B. Viral gene level correlation (viral gene expression vs fraction of immune cells)

**Figure 8:**
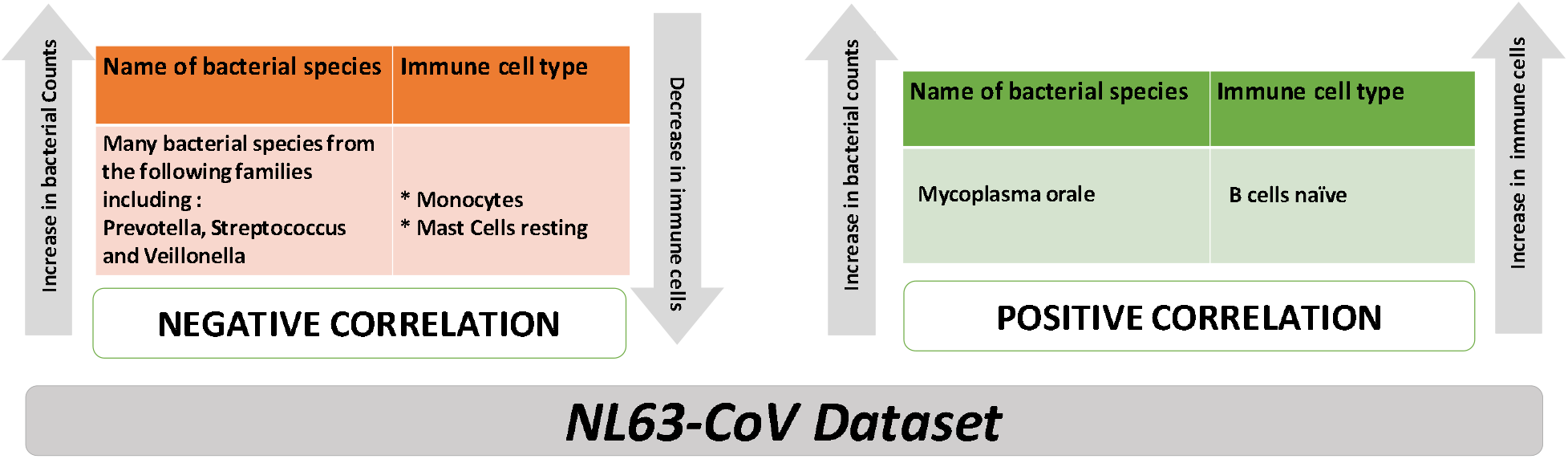
Summary of correlation analysis between bacterial abundance with immunological cell types for the NL63-CoV dataset.

**Figure 9.**
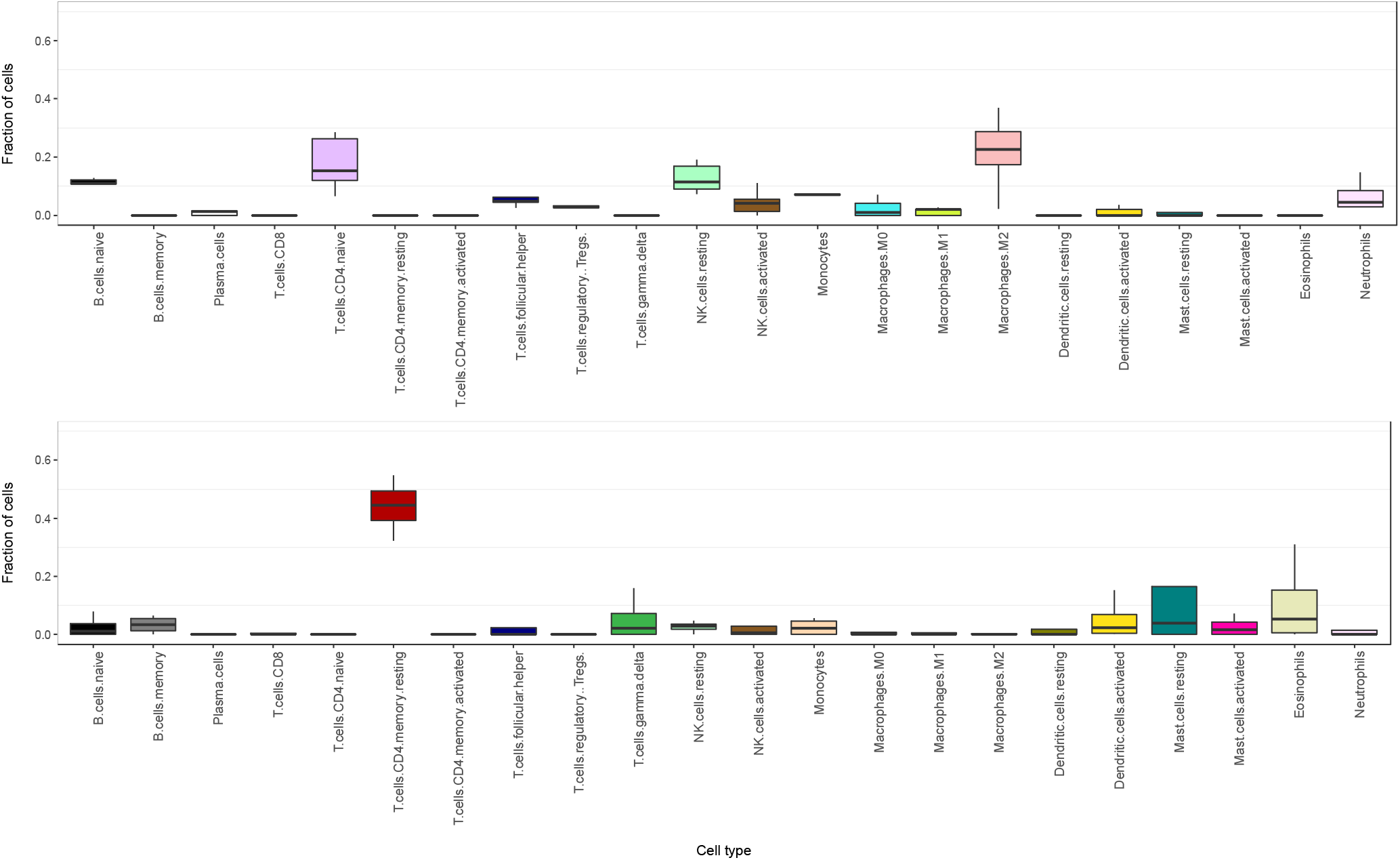
Box plot for 22 types of immune cells in (a) 5 patients from the SARS-COV-2 dataset and (b) 4 patients from the NL63-COV-Hiseq dataset. (Sample 1 from this cohort was ignored for this box plot due to low coverage)

We saw from our analysis of the SARS-CoV-2 dataset that Macrophages M2 are positively correlated with viral load (Figure 4A). Macrophages are special immune cells that detect, ingest and destroy target cells. It works by stimulating the immune system through M1 macrophages, and can also encourage tissue repair with the help of M1 macrophages [41, 42]. The macrophages located in the lungs are referred to as *alveolar macrophages*. If these macrophages are excessively activated, they could create a ‘cytokine storm’ which leads to release of pro-inflammatory factors such as interleukins, and hyper-inflammation; and causes immune cells to attack the organs in the body [43, 44]. Such a cytokine storm was commonly found in severe COVID-19 patients with Acute respiratory distress syndrome (ARDS) [44]. Recent publications have found that Macrophages M1 in the lung amplify and spread the COVID-19 viral infection, while Macrophages M2 degrade the SARS-CoV-2 virus and limit the spread of infection [45]. Scientists have been exploring treatments that target the pathways in order to lessen the cytokine storm effect in severe COVID-19 disease and some clinical trials are now underway [46].

Scientists are also researching ‘autoantibodies’ and their possible link to the long term side effects of Long COVID [47]. Autoantibodies are antibodies i.e. immune proteins that target a person’s own cells and organs, and causes autoimmune diseases such as Lupus [48]. For some patients that recovered from COVID-19, the immune profiled reverted back to normal, but for others researchers found many immune cells altered many months after recovery from COVID-19 when patients no longer test positive for the SARS-CoV-2 virus [49–52]. Liu et al [49] found depletion of NK cells and NK-like cells in patients recovering from COVID-19 [49]. The depletion of NK cells was also seen in our correlation results (Figure 4A). In other published articles, *Effector B and T cells* were found elevated, and was found associated with the long term side effects of the disease [50, 53–55]. These studies indicate that observing and mediating immune response, including T cell responses could be critical in the treatment of Long COVID.

Correlation analysis of the three types of measurements in the *SARS-CoV-2 dataset* has showed significant correlation between viral load as well as level of specific viral gene expression with the fractions of immune cells present in lung lavage as well as with abundance of major fractions of lung microbiome.

The role of NK cells is to provide quick response to virus-infected cells [56]. Our Correlation analysis showed that viral load of the SARS-CoV-2 genome was inversely correlated with natural killer (NK) cells activation (Figure 4A). In other words, the NK cells were inactivated in our analysis of the *SARS-CoV-2 dataset* (Figure 4A). Jewett et al [57] also found a correlation between reduction of NK cells and severity of disease [57]. Such depletion pattern of NK cells was also observed in patients affected with SARS virus [58]. NK cells are one of the most abundant types of lymphocytes in the lung, and could offer an interesting venue for COVID-19 treatments. Clinical trials are currently underway (NCT04634370) and scientists are now exploring immunotherapy and enhancement based treatments using NK cells to combat disease progression and severity of COVID-19 [57, 59–61].

Bacterial co-infections were not very frequent, but more commonly found in critically ill COVID-19 patients [62–64]. Bacterial pathogens found in the lungs of COVID-19 patients included *Enterobacter species, Pseudomonas, Streptococcus pneumoniae* were also found in our results (Figure 6). The *Enterobacteriaceae* species was found to be resistant to antibiotics in some CVID-19 patients [63]. Although not common, such infections in COVID-19 patients were complex to treat since it was not easy to distinguish the bacterial co-infections from viral infections of the respiratory tract. Our correlations results also show a high correlation of these bacteria with immune markers including Eosinophils and activated NK cells (Figure 5). This matches findings from Mason et al [65] who recommended studying inflammatory markers including Lymphocytes (such as NK cells, T cells, and B cells), and Neutrophils to distinguish the bacterial co-infections from viral infections of the respiratory tract [65].

### Immune landscape and correlation analysis in the *NL63-CoV* dataset

In our analysis of the NL63-CoV dataset, we saw the dominance of mast cells and CD4 memory resting T cells (Figure 9). Richards et al [54] examined circulating T cells in human endemic coronaviruses (hCoV) including HCoV-229E, NL63-CoV, OC43 and HKU1; and theorized that the *memory CD4 T-cells* found in patients exposed to infection from these endemic strains could also influence the immune response to SARS-CoV-2 infection and vaccination [53, 54].

### Findings from analyses on both *SARS-CoV-2* and *NL63-CoV* datasets

In a normal cell, T cells including CD4 and CD8 work to identify antigens from foreign pathogens. When such an event happens, the T cells differentiates into short lived *effector T cells* that work to control the foreign pathogens. When this task is done, the effector T cells are lost, but the preserved in the memory T-cells for long term immune response [66]. In published articles, *Effector B and T cells* were found elevated, and was found associated with the long term side effects of the disease [50, 53–55]. From the analysis on the *NL63-CoV dataset*, we saw that elevated *memory CD4 T-cells* could help in the long term immune response to the SARS-CoV-2 infection [53, 54]. This finding indicates that a treatment or vaccine that could potentially mediate the T-cell response to produce effective *memory CD4 T-cells;* and controlling the levels of *effector T-cell activity* may not only provide immunity from the SARS-CoV-2 infection, but also help prevent the adverse side effects in Long COVID in patients affected by this virus [66, 67].

Shaath et al [68] examined the blood samples from COVID-19 patients admitted to the hospital intensive care unit (ICU) vs those not in ICU and found several mRNA based markers associated with severity of the COVID-19 disease. The authors found several pathways related to NK cells and interferon signaling to be downregulated in the ICU COVID-19 patients. The authors recommended restoration of NK cells, and mediation of interferon-gamma as potential therapeutic options [68]. This work was done on blood samples and hence their findings are valid at a systemic immune response. This compliments our findings well which are about immune response at local tissue level.

### Relevance of machine learning tools and algorithms

As COVID-19 cases continue to rise around the world, researchers are harnessing the computational power of machine learning and artificial intelligence (AI) tools to not only create prediction and diagnostic tools for COVID-19 [69, 70] but also improve outcomes [71]. In this paper, we described the application of machine learning tools to process the raw sequencing data generated by NGS technology, and also explore the immune, viral and bacterial landscape of the *SARS-CoV-2 and NL63-CoV datasets*.

While traditional laboratory techniques allow direct detecting of immune cells in the blood, it is more difficult to do for other types of tissues where immune cells can be detected using a technique called flow cytometry or image cytometry methods. But both of these techniques are difficult and labor- and time-consuming.

RNA-sequencing (RNA-seq) technology in conjunction with the application of machine learning based virtual flow cytometry tools could be considered a potential alternative. Such an in-silico process would enable researchers to not only estimate the immune cell environment, but also work towards new hypothesis and therapies that would mediate appropriate immune response using T cells for long term immunity; and help to minimize adverse side effects from the SARS-CoV-2 infection [72, 73].

## CONCLUSION

In this paper, we applied multiple machine learning tools to NGS data analysis of lung tissue samples from COVID-19 patients. We explored the SARS-CoV-2 genome and compared it with other endemic coronavirus NL63 genome. Finally, we explored the immunological landscape of lung microenvironment from the *SARS-Cov-2 and* nasopharyngeal microenvironment from the *NL63-CoV datasets.*

Our exploratory study has provided novel insights into complex regulatory signaling interactions and correlative patterns between the viral infection, inhibition of innate and adaptive immune response as well as microbiome landscape of the lung tissue. Many of our findings from the analysis of the immune landscape of these two datasets, along with correlation analysis have been corroborated in published literature proving that the study of immune system warrants further analysis and exploration.

The study of how the SARS-CoV-2 virus interacts with the immune system; and comparing and contrasting the immune system in patients affected by endemic viruses could offer important insights into protection against SARS◻CoV◻2; and shed light on new therapies to combat severe COVID-19 disease.

These initial findings on small group of samples could provide better understanding of the diverse dynamics of immune response and the side effects of the SARS-CoV-2 infection but require further validation on a larger cohort of samples.

## LIST OF SUPPLEMENTARY FILES

- Supplementary File 1A: shows the estimated copy number of viral genomes detected in lung lavage samples of the SARS-CoV-2 dataset obtained using the viGEN pipeline.
- Supplementary File 1B: CENTRIFUGE metagenomics pipeline results on the SARS-CoV-2 dataset
- Supplementary File 1C: shows the estimated viral level of viral gene expression counts
- Supplementary File 2A: correlation between genome level abundance and immunological data for the SARS-CoV-2 dataset
- Supplementary File 2B: : correlation between gene level abundance and immunological data for the SARS-CoV-2 dataset
- Supplementary File 2C: Correlation between bacterial abundance with immunological cell types for the SARS-CoV-2 dataset
- Supplementary File 2D: Correlation between genomic viral load with bacterial abundance for the SARS-CoV-2 dataset
- Supplementary File 3A: shows the estimated viral genome copy numbers in 5 pediatric patients from the NL63-CoV dataset obtained using the viGEN pipeline
- Supplementary File 3B: CENTRIFUGE metagenomics pipeline results on the NL63-CoV dataset
- Supplementary File 3C shows the NL63 viral gene/CDS counts in the nasopharyngeal microenvironment of the NL63-CoV dataset
- Supplementary File 4 shows the microbiome profile of pediatric patients from the nasopharyngeal microenvironment in the NL63-CoV dataset represented as a Sankey diagram visualization of the bacterial species.
- Supplementary File 5A: correlation between genome level abundance and immunological data for the NL63-CoV dataset
- Supplementary File 5B: : correlation between gene level abundance and immunological data for the NL63-CoV dataset
- Supplementary File 5C: Correlation between bacterial abundance with immunological cell types for the NL63-CoV dataset
- Supplementary File 5D: Correlation between genomic viral load with bacterial abundance for the NL63-CoV dataset

## LIST OF ABBREVIATIONS

COVID-19: The coronavirus disease 2019
SARS-CoV-2: SARS coronavirus 2
SARS-CoV: SARS coronavirus
IFN: Type I interferons
NCATS: NIH National Center for Advancing Translational Sciences
EHRs: Rlectronic health records
WSI: Whole slide images
PASC: Post-acute sequelae of SARS-CoV-2 infection
HCoV-NL63: NL63 coronavirus
hCoV: Human endemic coronaviruses
NGS: Next generation sequencing
SIRS: Severe acute respiratory infection
SIMD: Single-instruction multiple-data
SB: Seven Bridges
CGC: Cancer Genomics Cloud Platform
ν-SVR: Nu-linear support vector regression
SVM: Support vector machine
BWT: Burrows-Wheeler transform
FM: Ferragina-Manzini index
NK: Natural Killer cells
AI: Srtificial intelligence
RNA-seq: RNA-sequencing

## CONFLICT OF INTEREST

All authors declare no conflict of interest.

## AUTHOR CONTRIBUTIONS

YG designed the analysis and study and performed interpretation of the results. KB performed the analysis. KB, YG and SM wrote and edited the paper.

## ACKNOWLEDGMENTS

The authors would like to acknowledge the CGC Seven Bridges team for enabling some of the high-throughput analysis using the CENTRIGUE metagenomic tool

## DATA AVAILABILITY STATEMENT

The datasets used in this manuscript are available online.

### *SARS-CoV-2* Dataset

We downloaded RNA-seq data from 5 patients affected with the SARS COV 2 virus from the NCBI SRA data repository PRJNA605983 (SRP249613) [22, 23]. These 5 patients were from the early stage of the Wuhan seafood market pneumonia virus outbreak in China. The downloaded data were raw sequences in the form of .FASTQ files. Total RNA was extracted from bronchoalveolar lavage fluid and then next generation sequencing (NGS) was performed using the Illumina platform. Out of the 5 samples, 4 were profiled on the Illumina HiSeq 3000 platform, and one on the Illumina HiSeq 1000 platform.

### Human NL63 coronavirus dataset (referred to as the *NL63-CoV* dataset in this paper)

We found a public dataset was from 5 pediatric patients with severe lower respiratory infection by NL63 coronavirus with deep sequencing data performed on Illumina HiSeq platform. The downloaded data were raw sequences in the form of .FASTQ files obtained from NCBI PRJA601331 [24, 25].

## Notes

### Competing Interest Statement

The authors have declared no competing interest.

