## Supplementary figures and images for "Computational genomic analysis of the lung tissue microenvironment in COVID-19 patients"

### Supplementary File 4

#### Sample1

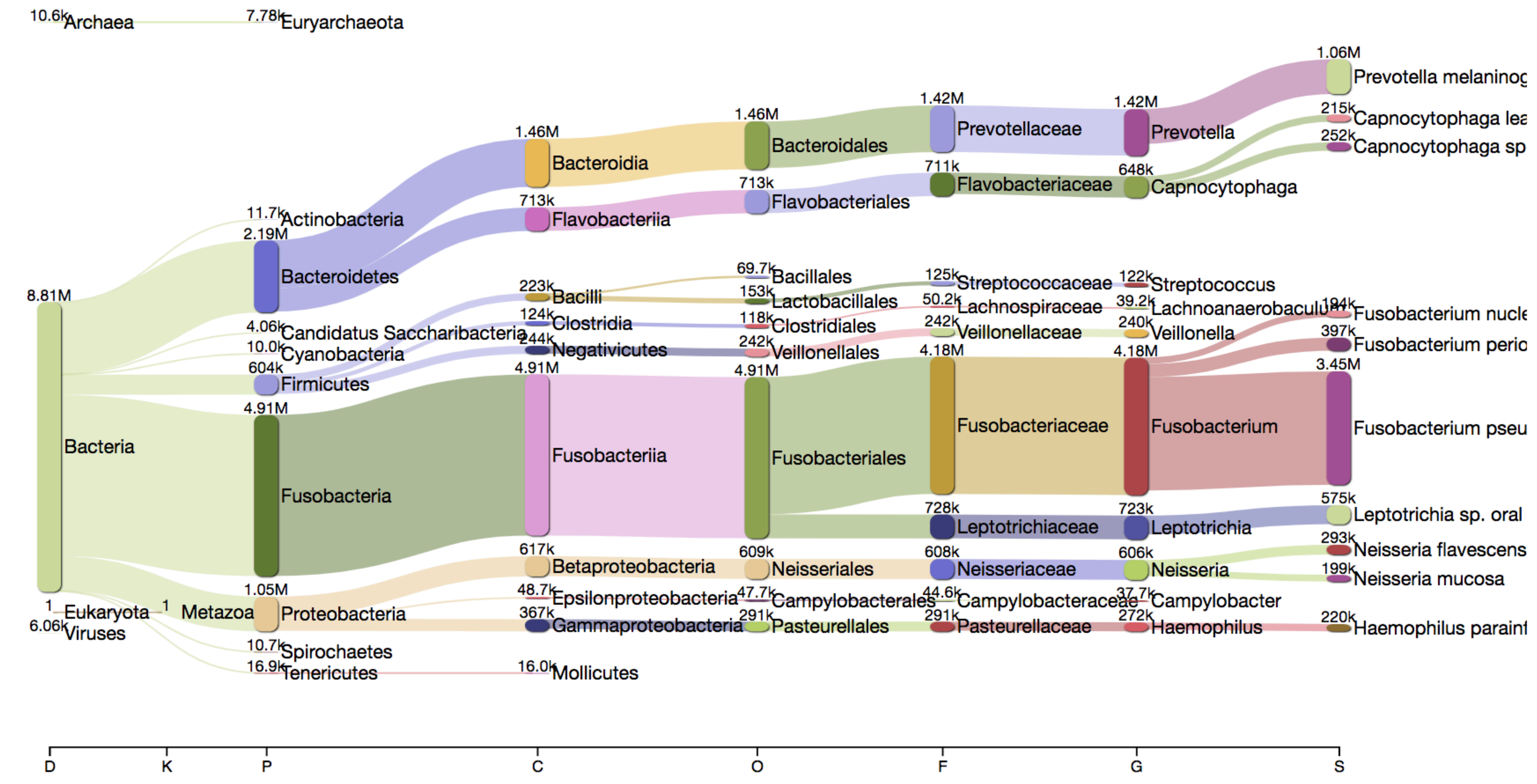


#### Sample2

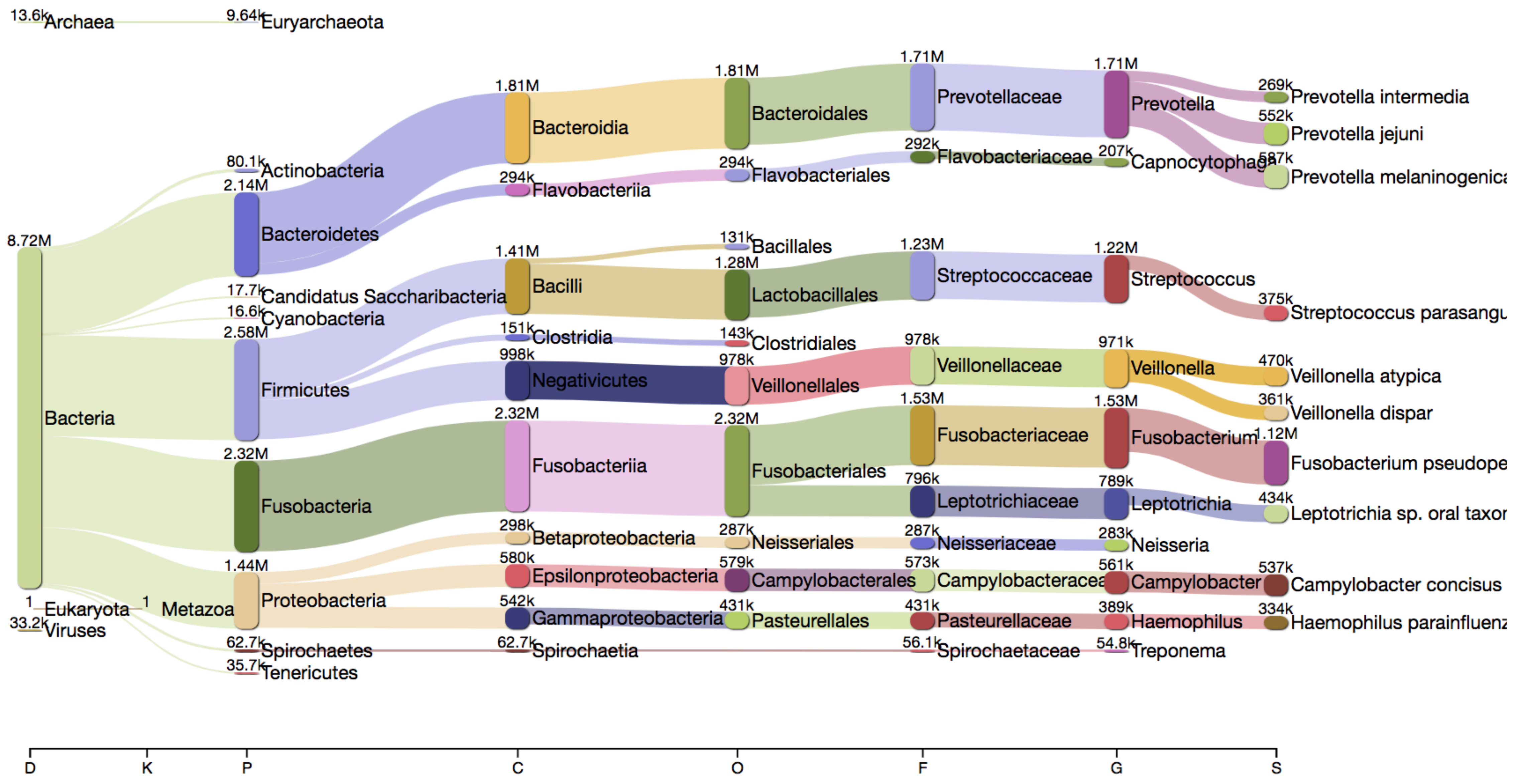


#### Sample3

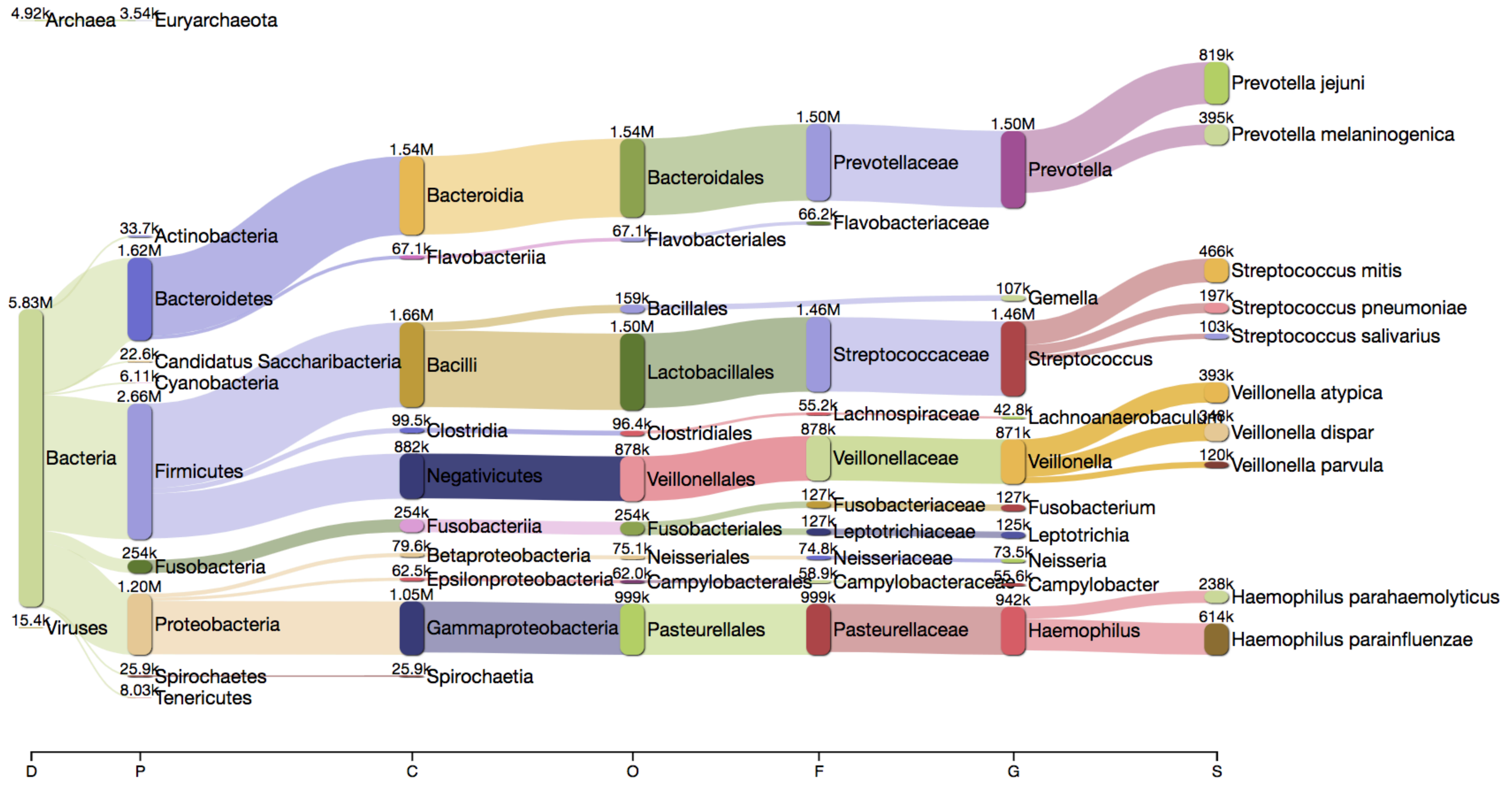


#### Sample4

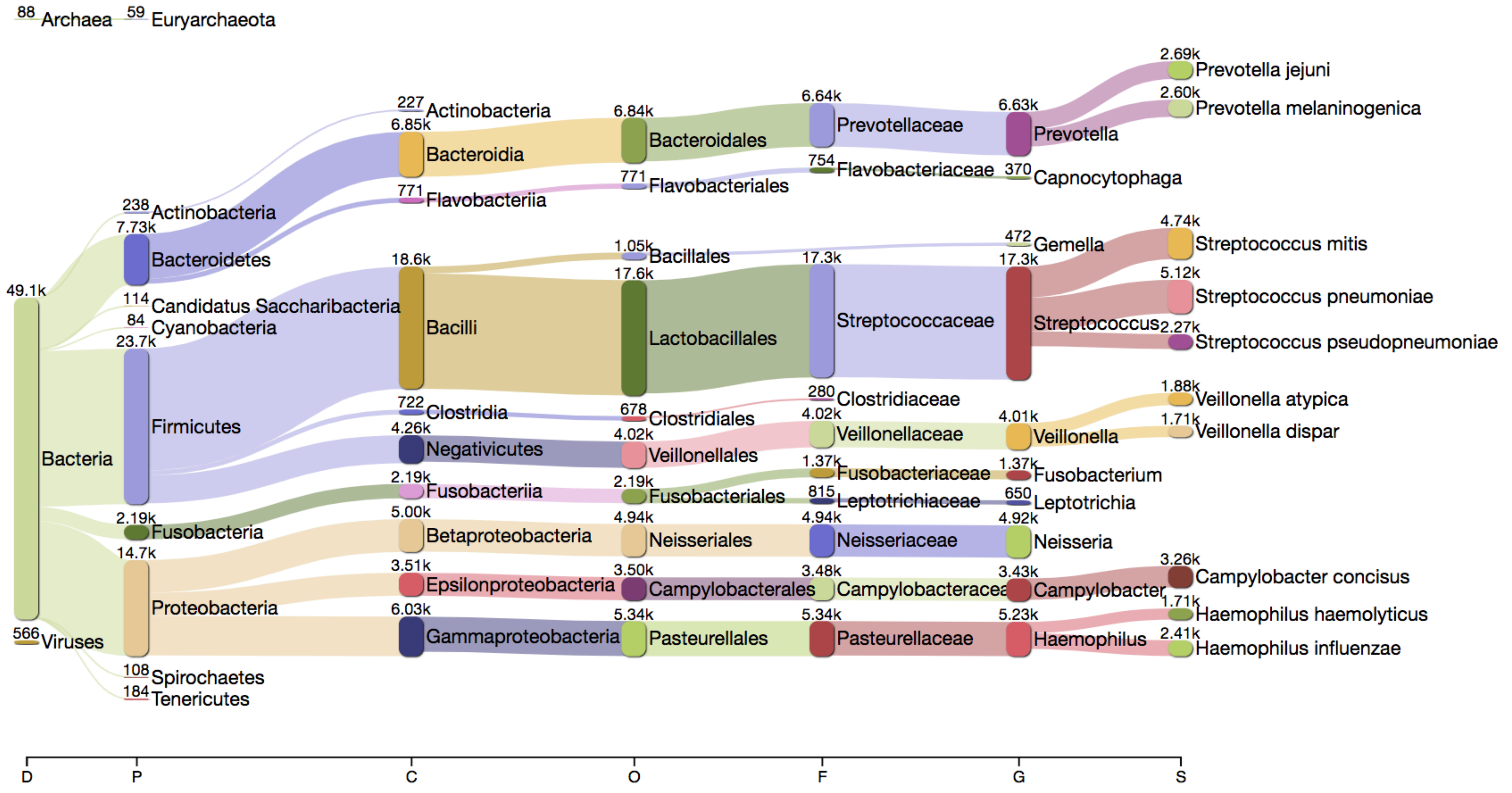


#### Sample5

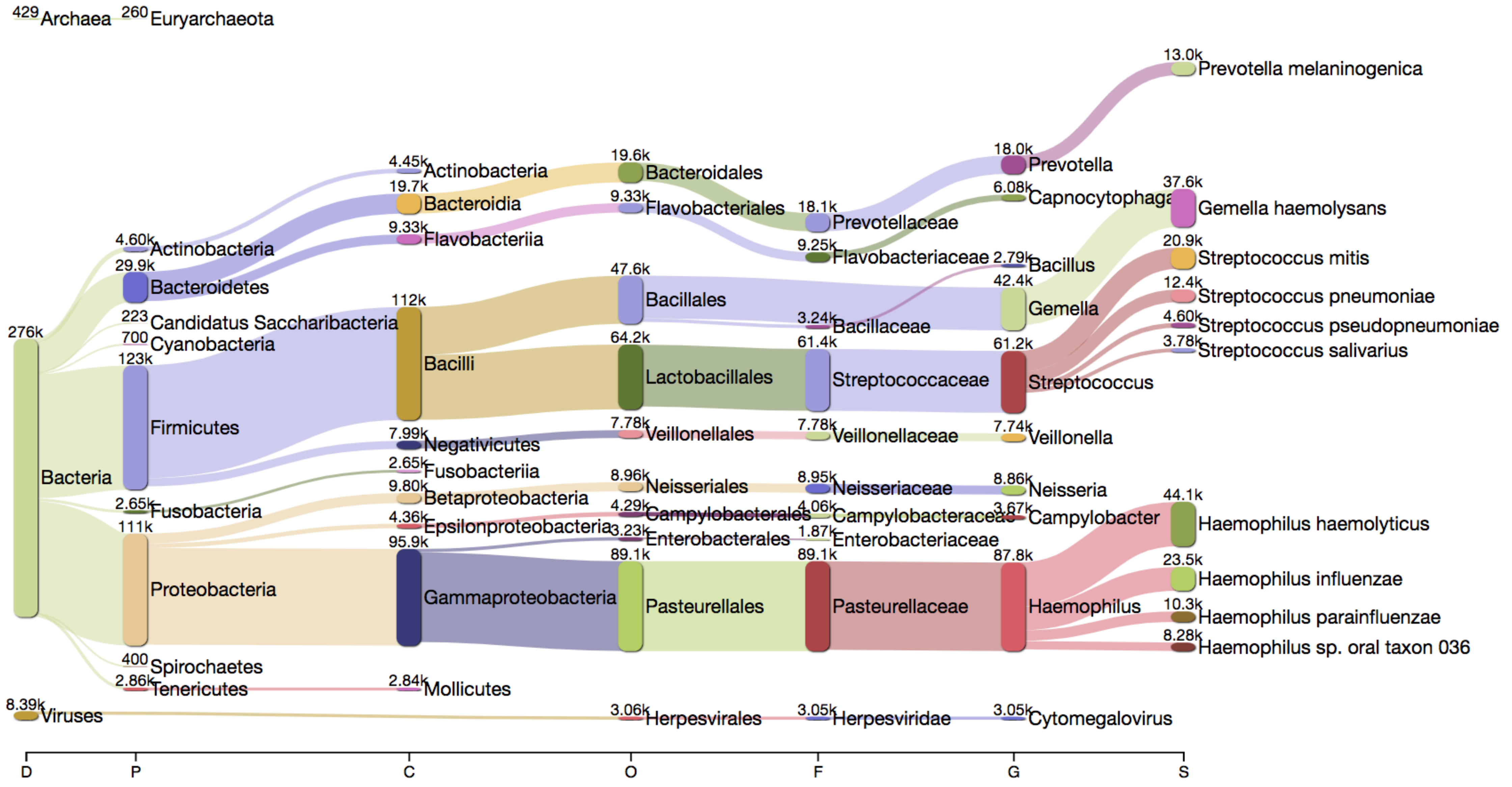
